# Insulator Activities of Nucleosome-Excluding DNA Sequences Without Bound Chromatin Looping Proteins

**DOI:** 10.1101/511600

**Authors:** Yuki Matsushima, Naoaki Sakamoto, Akinori Awazu

## Abstract

Chromosomes consist of various domains with different transcriptional activities separated by chromatin boundary sequences such as insulator sequences. Recent studies suggested that CTCF or other chromatin loop-forming protein binding sequences represented typical insulators. Alternatively, some long nucleosome-excluding DNA sequences were also reported to exhibit insulator activities in yeast and sea urchin chromosomes although specific binding of loop-forming proteins were not expected for them. However, the mechanism of the insulator activities of these sequences and the possibilities of similar insulators existing in other organisms remained unclear. In this study, we first constructed and performed simulations of a coarse-grained chromatin model containing nucleosome-rich and nucleosome-excluding DNA regions. We found that a long nucleosome-excluding region between two nucleosome-rich regions could markedly hinder the associations of two neighboring chromatin regions owing to the stronger long-term-averaged rigidity of the nucleosome-excluding region compared to that of nucleosome-rich regions. Subsequent analysis of the genome wide nucleosome positioning, protein binding, and DNA rigidity in human cells revealed that some nucleosome-excluding rigid DNA sequences without bound chromatin looping proteins could exhibit insulator activities, functioning as chromatin boundaries in various regions of human chromosomes.

## INTRODUCTION

Eukaryotic chromosomes contain various hierarchical chromatin structures such as topologically associated domains and A/B compartments;^1–8^ in addition, transcriptional activities are variously regulated among individual chromatin domains. Recent studies have suggested that the DNA sequences on which CTCF can bind represent typical sequences comprising the boundary domains among such chromatin domains in various organisms.^9–20^ Such sequences have been recognized as typical insulator sequences that can insulate the influences of enhancer associations and heterochromatin extension from neighboring chromatin domains by making contact with each other to form a loop from each chromatin domain. In human and yeast cells, RNA polymerase III-transcribed sequences were also suggested to serve as insulator sequences with similar mechanisms as CTCF binding sequences.^19–28^ Moreover, various protein binding sequences that form chromatin loops by contacting each other and also exhibit insulator activities have been found in various invertebrates.^19,20,29–34^

To precisely regulate gene expression, each enhancer associates with promoters only within the same chromatin domain, with each domain divided by insulators. To explain the mechanisms underlying such physically nontrivial limitation of enhancer motion, various models of chromatin dynamics and enhancer-promoter interactions such as the tracking model for DNA binding proteins, looping model based on enhancer-promoter random collision or enhancer scanning, and the topological domain model based on enhancer-promoter random collision have been proposed.^35–41^ Notably, recent experiments and mathematical models suggested that enhancer-promoter interactions could be sufficiently reproduced by a topological domain model wherein the topological domains were assumed to be formed by chromatin loop-forming protein binding to insulator sequences.^40,42–44^ In particular, the promoter could associate with the enhancer in the same chromatin domain more frequently than with that in the neighboring chromatin domain even if the genomic distances from both enhancers to the promoter were identical.^42^

To function as chromatin domain boundaries, it is necessary for the insulator sequences to associate with other insulator sequences via their binding proteins, such as the CTCF-cohesin complex.^9–20^ Alternatively, some nucleosome-excluding sequences such as (A)_n_ or (CCGNN)_n_ repeat sequences and the core 182 bp region of *Ars* insulator sequences (ArsInsC) found in sea urchin genome have been shown to independently exhibit insulator activities.^45,46^ For these sequences, the specific binding of proteins such as CTCF to form chromatin loops was not identified. Moreover, the insulator activities of CTCF binding sequences are highly dependent on their intra-genomic direction^5,10,11,16^ whereas those of nucleosome-excluding sequences are robust.^45,46^ This indicates the possibility that such nucleosome-excluding sequences may exhibit insulator activities without the formation of chromatin loops; herein these are therefore termed as nucleosome-excluding non-looping insulator sequences (NENLIS). However, the physical mechanism of the NENLIS-mediated insulator activities remained unknown.

Furthermore, the *Ars* insulator sequence was demonstrated as showing insulator activities not only in sea urchin cells but also in cells of various organisms including human and plants.^47–50^ This suggested that NENLIS might be distributed in various chromosomal regions and exhibit insulator activities across different organisms. However, despite the exhaustive epigenome and 3-D chromosome structure data obtained via Hi-C experiments from various research projects, little progress has been made regarding the genome-wide analysis of such sequences.^51–54^

In this study, we constructed a coarse-grained model of a chromatin chain consisting of a nucleosome-rich region and NENLIS to propose possible scenarios for the exhibition of insulator activities by long nucleosome-excluding DNA sequences. We also performed an exhaustive search of the expected NENLIS through the analysis of genome wide micrococcal nuclease (MNase) data of human cells, with a focus on the contact probabilities among loci around NENLIS as ascertained through analysis of Hi-C data, to evaluate their insulator activities.

## MATERIALS AND METHODS

### Coarse-Grained Model of a Chromatin Chain containing a Nucleosome-rich Region and NENLIS

The role of insulator sequences is as a hindrance to the interactions between two chromatin regions belonging to different chromatin domains. Thus, to simulate the insulating behaviors and evaluate the activities of NENLIS, we constructed a coarse-grained model of a chromatin chain containing nucleosome rich regions and NENLIS.

We focused on the steady state behaviors of the chromatin chain over a long time scale wherein each chromatin (nucleosome) remodeling event could be regarded as noise. In this case, we could regard the chromatin chain as a chain with inhomogeneous rigidity (bending elasticity), where the regions corresponding to NENLIS were much rigid compared to those corresponding to nucleosome-rich regions.

In nucleosome rich regions, the nucleosome positions on DNA sequences often shift along the sequences as a result of the thermal fluctuations and chromatin remodeling factor activities^55–65^. In particular, when the position of a histone octamer on DNA sequence shifts 1 bp, the angle of the helical axis of DNA wrapped around the nucleosome shifts approximately 34 degrees, as the helical period of double-strand DNA is approximately 10.5 bp. Therefore, the positional relationship among nearby nucleosomes can change largely because of the small shifts of a histone octamer along the DNA sequence. For example, in the case of (A)_750_ sequence that contains four nucleosomes, the distance between the nucleosomes furthest up- and downstream changes between 10–35 nm by the 1–20 bp shifts of the position of the second-most upstream nucleosome along the DNA sequence (Figure 1). Here, this evaluation was performed by the following procedures using online Interactive Chromatin Modeling (ICM) (http://dna.engr.latech.edu/icm-du/index.php):^66^

**Figure 1.**
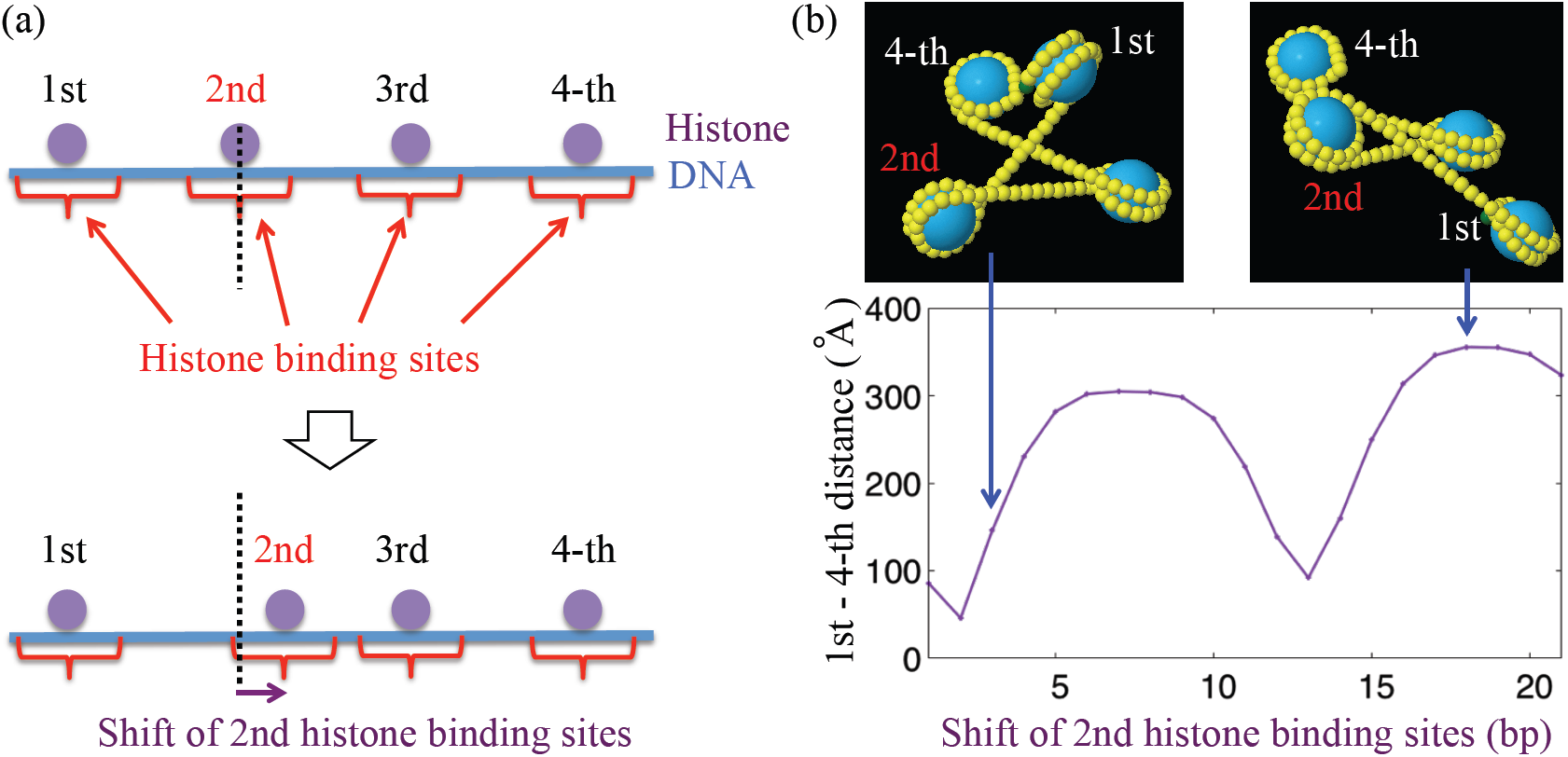
Histone binding site-dependent variation of local chromatin shapes. (a) Illustration of shift of histone binding site. (b) Typical local chromatin structures that vary with the difference of 2nd histone binding site, where blue and yellow particles indicate histone octamer and 5 bp DNA (upper), and the physical distance between the 1st and 4th nucleosomes as a function of the shift of 2nd histone binding sites (lower). Local chromatin structure markedly changes according to histone binding site.

I. “K = MD-B.dat” and “X_DNA_ = MD-B.par” were chosen as the DNA model and “X_nuc_ = 01kx5.min” was employed as the nucleosome model.
II. The nucleosome positions were given as 1st, (201+n)th, 401st, and 601st bp, where four nucleosomes were wrapped respectively by the 1st to 147th, (201+n) to [(201+n)+147]th, 401st to 547st, and 601st to 747th bp.
III. For n = 0 to 20, the distances between the 1st and 4th nucleosomes were measured. Similar inter-nucleosome position-dependent chromatin structure variations have also been observed in *in vitro* experiments and simulations using mathematical models.^55–65,67,68^

Accordingly, the nucleosome-rich region could be regarded as a flexible chain if we focused on sufficiently longer time-scale behaviors of the chromatin chain than the time interval between which the majority of associations of chromatin remodeling factors to each nucleosome occur. Moreover, based on its elasticity, NENLIS deforms little from the basic double strand DNA structure during such longer time-scale behaviors although it exhibits fast small fluctuations resulting from thermal noise. This indicated that we could model the nucleosome-rich region and NENLIS as flexible and rigid chains, respectively, if we focused on the statistical properties of sufficiently longer-time behaviors of chromatin regions with NENLIS.

### Implementation of the Coarse-grained Chromatin Model

Based on the abovementioned considerations, we constructed the coarse-grained chromatin models using the bead-spring model with excluded volume effects and two-body nucleosome interactions as follows (Figure 2). The chains consisted of *N* spherical particles with diameter *d*, and the *i*-th particles (*i* = 1, 2, … *N*) were coupled with (*i*-1)th and (*i*+1)th particles via the spring with the natural length = *d*. Here, we assumed *d* = 11 nm, which is similar to the length of approximately 32 bp DNA and the diameter of the spherical region containing a single nucleosome (147 bp DNA wrapping each histone octamer) with euchromatin linker DNA (approximately 30 bp).^69^ For simplicity, we described the nucleosome-rich region and NENLIS using the particles with the same *d*, wherein we assumed that each particle in the nucleosome-rich region contains approximately 180 bp DNA and that in NENLIS contains approximately 32 bp DNA.

**Figure 2.**
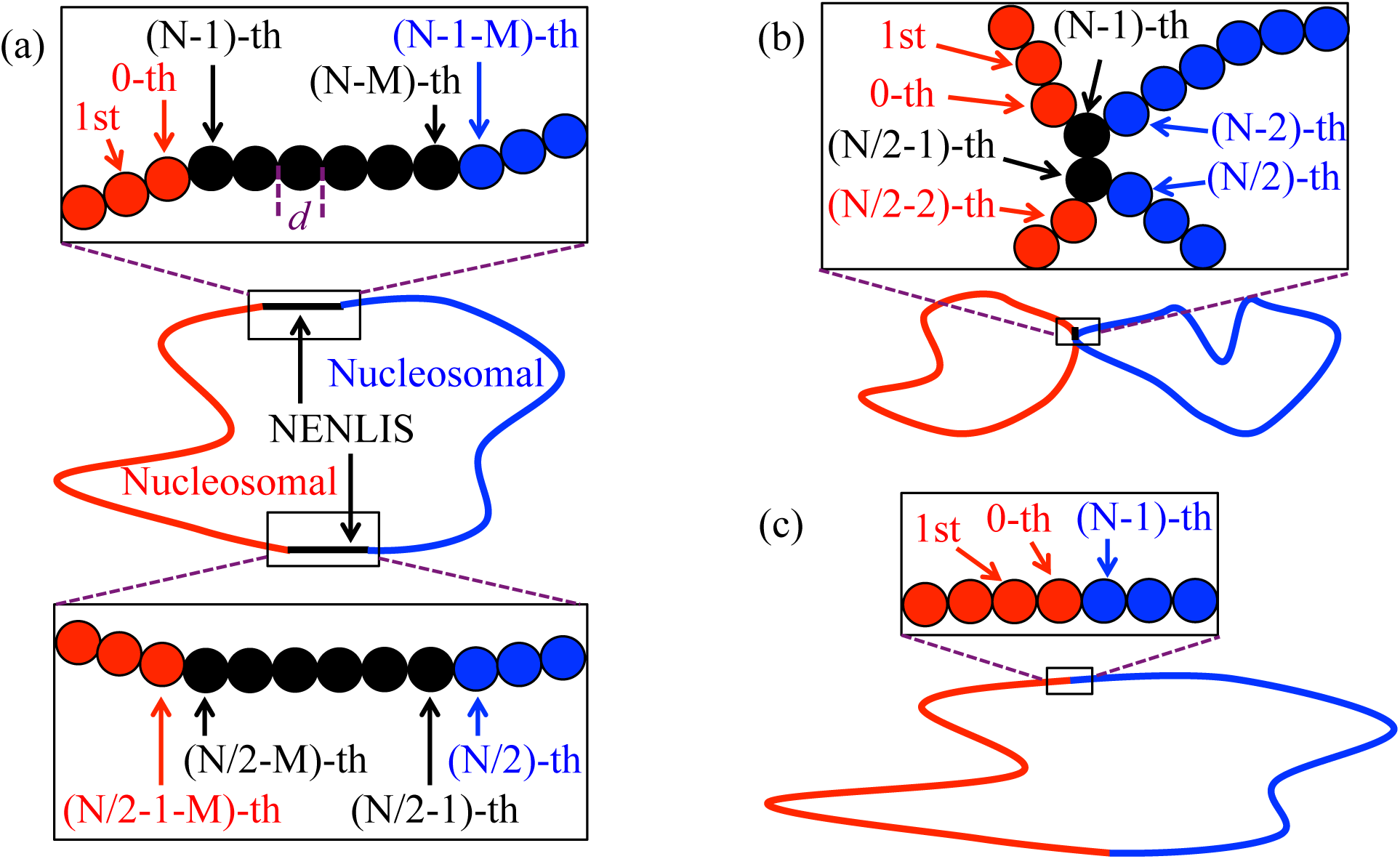
Illustrations of particle chain models of chromatin. (a) Illustration of the NENLIS-ring, where red and blue particles represent nucleosome-rich (nucleosomal) regions and black particles represent NENLIS. The diameter of each particle is *d* and each index represents the particle index. (b) Illustration of Two-coupled-rings where the two black particles represent a permanently connected chromatin region. (c) Illustration of the control ring.

We assumed that the motion of the *i*-th particle obeys the following Langevin equation:

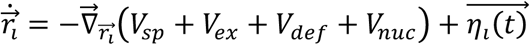

where 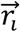 indicates the 3-D coordinate of the *i*-th particle, *V*_*sp*_, *V*_*ex*_, *V*_*def*_, and *V*_*nuc*_ indicate the potential of forces working on particles, and 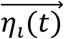 indicates the Gaussian white noise.

The potential *V*_*sp*_ indicates the elastic potential of the spring connecting two particles given as:

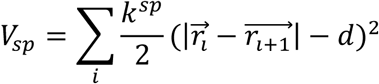

where *k*^*sp*^is the spring constant. *V*_*ex*_ indicates the potential of the excluded volume between two particles given as:

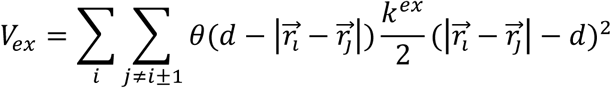

where *k*^*ex*^ is the elastic constant when two particles contact with each other, and *θ* indicates the Heaviside step function.

To implement the differences of the rigidity between the nucleosome-rich region and NENLIS region in the chromatin chain, the potential *V*_*def*_ was considered as:

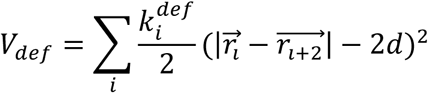

where 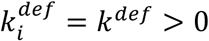 if the *i*-th, (*i*+1)th, and (*i*+2)th particles belong to NENLIS regions, and 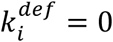 otherwise. The combination of *V*_*def*_ and *V*_*sp*_ provided the differences of the rigidity between the nucleosome-rich region and NENLIS region; the particle chains corresponding to nucleosome-rich regions are flexible whereas those corresponding to NENLIS regions tend to remain straight.

The potential *V*_*nuc*_ indicates that the attractive interaction potential among particles belongs to the nucleosome-rich regions. Recent studies suggested that the attractive forces between two nucleosomes occur through the interaction among their histone tails and various histone tail binding proteins.^69–76^ Based on these observations and simulations of their model Mukhopadhyay et al.^42^ concluded that each nucleosome tends to exhibit attractive interaction with only one other nucleosome. Accordingly, we assumed the attractive force is active between two particles if they belong to the nucleosome-rich regions and form a pair. Then, *V*_*nuc*_ is given as:

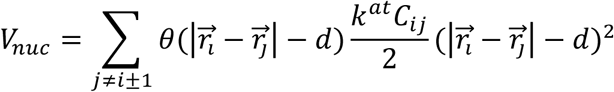

where *k*^*at*^ indicates the coefficient of attractive potential. Here, *C*_*ij*_ = 1 when the *i*-th and *j*-th particles belong to the nucleosome-rich region and form a pair, and *C*_*ij*_ = 0 otherwise. We also assumed that *C*_*ij*_ changes temporally as in the following rules:

i. The changes *C*_*ij*_ = 0 → 1 and *C*_*ji*_ = 0 → 1 occur when *C*_*ii*’_ = 0 for all *i’*, *C*_*jj*’_ = 0 for all *j’*, *j* ≠ *i* ± 1, and 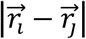 becomes < *d*.
ii. The changes *C*_*ij*_ = 1 → 0 and *C*_*ji*_ = 0 → 1 occur when *C*_*ij*_ = 1 and 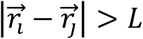. In the present study, the case of *L* = 1.01*d* was considered; however, we confirmed the results for the cases of *L* = 1.005*d* and *L* = 1.02*d* were qualitatively similar to *L* = 1.01*d* (see Figure S1).

The Gaussian white noise 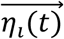 obeys 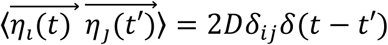. Here, we assumed that *D* is independent of the temperature of the system as the origin of this noise includes the influences of various proteins such as chromatin remodeling factors and RNA polymerase on the chromatin chain, along with energy consumption such as ATP hydrolysis.

### Evaluation of the Insulator Activities of NENLIS in the Coarse-grained Model

We performed long-term simulations of the coarse-grained chromatin chain model using the following parameter set: *k*^*sp*^*d*^2^/*D* = *k*^*ex*^*d*^2^/*D* = *k*^*def*^ *d*^2^/*D* = 200 ≫= 1, which were assumed to maintain the total particle volume of the chains and render each NENLIS region as a rigid straight chain. We considered the cases of *k*^*at*^*d*^2^/*D* = 200, 20, 2 and = 0 (*k*^*at*^ = 0), which indicate the cases of the presence and absence of two-body internucleosome attraction. In the following arguments, we focused on the long-term average of the steady state behaviors of the model. For simulations, we employed the particular values of parameters as *d* = 1, *D* = 50, *k*^*sp*^ = *k*^*ex*^ = *k*^*def*^ = 10000, and *k*^*at*^ = 10000, 1000, 100 or 0, that satisfies the abovementioned conditions; however, the arbitral values of them did not influence our focused properties.

To evaluate the insulator activities of NENLIS, we compared the behaviors of the following three models of chromatin rings consisting of N particles.

a. NENLIS-ring: One chromatin ring in which the nucleosome-rich regions were divided into two regions with the same length by two NENLIS regions with the same length; Nucleosome-rich regions were described by two sets of particles from the 0-th to (N/2-1-M)th and (N/2)th to (N-1-M)th particles, and NENLIS were described from the (N/2-M)th to (N/2-1)th and (N-M)th to (N-1)th particles (Figure 2a).
b. Two-coupled-rings: Two chromatin rings with the same chain length were formed by the division of one ring of a nucleosome-rich region by the eternal connection of two particles (these two particles were termed boundary particles and assumed to be always connected). In particular, one ring contains the particle set from the 0-th to (N/2-2)th particles, the other contains that from (N/2)th to (N-2)th particles, and the two rings were constructed by the eternal connection of the (N/2-1)th and (N-1)th particles (Figure 2b). Notably, this system is almost the same as that studied by Mukhopadhyay et al.^42^
c. Control-ring: One chromatin ring that contains only the nucleosome-rich region.

The insulator activity of NENLIS was evaluated by the comparison with contact maps (contact probability matrix) among particles belonging to nucleosome-rich regions obtained from the simulation of the three types of chromatin rings. The contact probabilities between the *m*-th and *n*-th particles belonging to nucleosome-rich regions in the cases of the NENLIS-ring, Two-coupled-rings, and Control-rings, 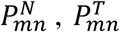 and 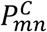 were defined by the probabilities exhibiting 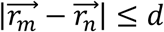 in the respective cases. We also measured relative contact probabilities 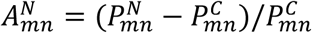 and 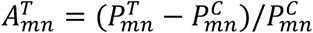 to evaluate the insulator activities in the NENLIS-ring and Two-coupled-rings where 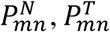, and 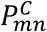 were measured under the same parameters as particles corresponding to the nucleosome-rich regions.

### Identification of NENLIS Candidates (NENLISc) in Human Cells

From the human genome sequence data (hg19), we identified candidate NENLIS sequences using genome-wide nucleosome positioning score, CTCF binding score, and PolIII binding score data from MNase and ChIP-Seq experiments of the ENCODE project.^52^ Herein, we focused on the data from the human GM12878 cell line.

Using MNase data (GEO: GSM920558),^77^ we first searched sequences > 150 bp at which the nucleosome scores were 0 for all loci. From these, we chose the sequences for which the fold changes over control of CTCF binding score (GEO: GSM935611) of all loci in each sequence and upper and lower 10 kbp sequences were < 1. Then, we further chose the sequences where the PolIII binding scores (GEO: GSM935316) of all loci in each sequence and upper and lower 10 kbp sequences were < 1.5. The resulting chosen sequences were not expected to constitute CTCF binding or PolIII transcribed sequences, or to form any chromatin loops. We next removed the sequences wherein the genomic distance between one of the sequence edges and the gene transcribed regions was < 500 bp as determined using human genome data (GENCODE Release 28 (mapped to GRCh37, hg19), https://www.gencodegenes.org/releases/28lift37.html) because such sequences might stimulate gene promoters or the promoter insulators.^78–83^ From the remaining sequences, we chose the physically rigid sequences as recently described NENLIS were expected to be rigid.^45,46^ Specifically, using the recently reported data set of the rigidities of the four nucleotide sequences,^84^ we defined the rigidity index of each sequence by averaging the rigidities of four sequential nucleotide series. The maximum, central, and minimum values of the rigidities in this data set were 1.9, 11.9, and 27.2. As the rigidity index of ArsInsC sequence was estimated as 14.7, we focused on sequences with rigidity indices > 15 as NENLIS candidate sequences (NENLISc). Through these procedures, we collected the NENLISc in GM12878 cells.

### Evaluation of the Insulator Activities of NENLISc in Human Cells

To evaluate the insulator activities (the ability to function as a chromatin domain boundary) of NENLISc, we measured the contact probabilities among loci belonging to the 10 kbp up- and downstream chromatin regions of NENLISc using Hi-C data of GM12878 cells with 1 kbp resolution (GEO: GSE63525). We defined 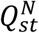 as the contact probability between the *s*-th and *t*-th loci around a NENLISc. Here, *s* and *t* were the relative positions from the focused NENLISc given as *s*, *t* = −10, −9, …, −2, −1, 1, 2, …,10 (kbp). We considered the case that Hi-C data provides the contact probability between two loci such that the relative positions from the central locus of the focused NENLISc to the loci are given as *s’* and *t’*, respectively. Then, 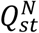 is given by the contact probability between the *s’*-th and *t’*-th loci wherein the sign of *s’* (*t’*) and *s* (*t*) are the same and |*s’*| (|*t*’|) is smaller than |*s*| (|*t*|) and larger than {|*s*| -1 kbp} ({|*t*| -1 kbp}). We also measured the contact probabilities between the *s*-th and *t*-th loci around 11,000 (5000 from each of 22 chromosomes) randomly chosen 200 bp sequences in a similar manner, and defined 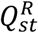 as the average contact probabilities around these 200 bp sequences. The relative contact probabilities are estimated by 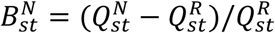.

Using 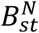, the insulator activities of each NENLISc were evaluated as follows. i) The average values of 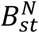 for regions with *s*, *t* < 0, *s*, *t* > 0, and *s* < 0, *t* > 0 (or *s* > 0, *t* < 0) were defined as 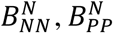, and 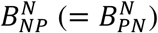, respectively. Here, 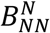, and 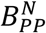 provide the average association strength among loci in the respective two chromatin regions whereas 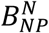 provides those among loci belonging to different chromatin regions divided by NENLISc. ii) We regarded the NENLISc as an insulator if 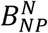 was significantly smaller than 0 with a P-value from t-test < 0.05.

## RESULTS AND DISCUSSION

### Relative Contact Probabilities and Expected Insulator Activities of the Coarse-grained Model

We performed simulations of the i) NENLIS-ring, ii) Two-coupled-rings, and iii) Control-rings. Here, we considered the cases that the NENLIS region in i) contained M = 6 particles where NENLIS was assumed as slightly shorter than 200 bp, similar to ArsInsC.^46^

The relative contact probabilities of NENLIS-ring 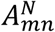 in the case of *N =* 112, 212, and 312 (approximately 18, 36, and 54 kbp) and those of Two-coupled-rings 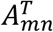 in the case of *N =* 100, 200, and 300 were measured (Figure 3). Here, the number of particles corresponding to the nucleosome-rich region was 100, 200, and 300 in both models. We found that 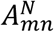 among the loci belonging to different nucleosome-rich regions in the NENLIS loop tended to be negative in the case of *N =* 112 (Figure 3a). This indicated that the associations among loci in different chromatin domains were specifically hindered compared to those among loci in the control ring owing to the rigidity of NENLIS; i.e., NENLIS provided a chromatin boundary as an insulator. For larger *N* such as *N* = 212 or 312, 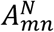 for the *m*-th and *n*-th particles, for which the distance to the NENLIS along the particle chain was smaller than the length of 50 particles, exhibited similar features to that for the case of *N* = 112 (Figure 3b and 3c). In contrast, the contact among *m*-th and *n*-th particles far from NENLIS could not be sufficiently hindered.

**Figure 3.**
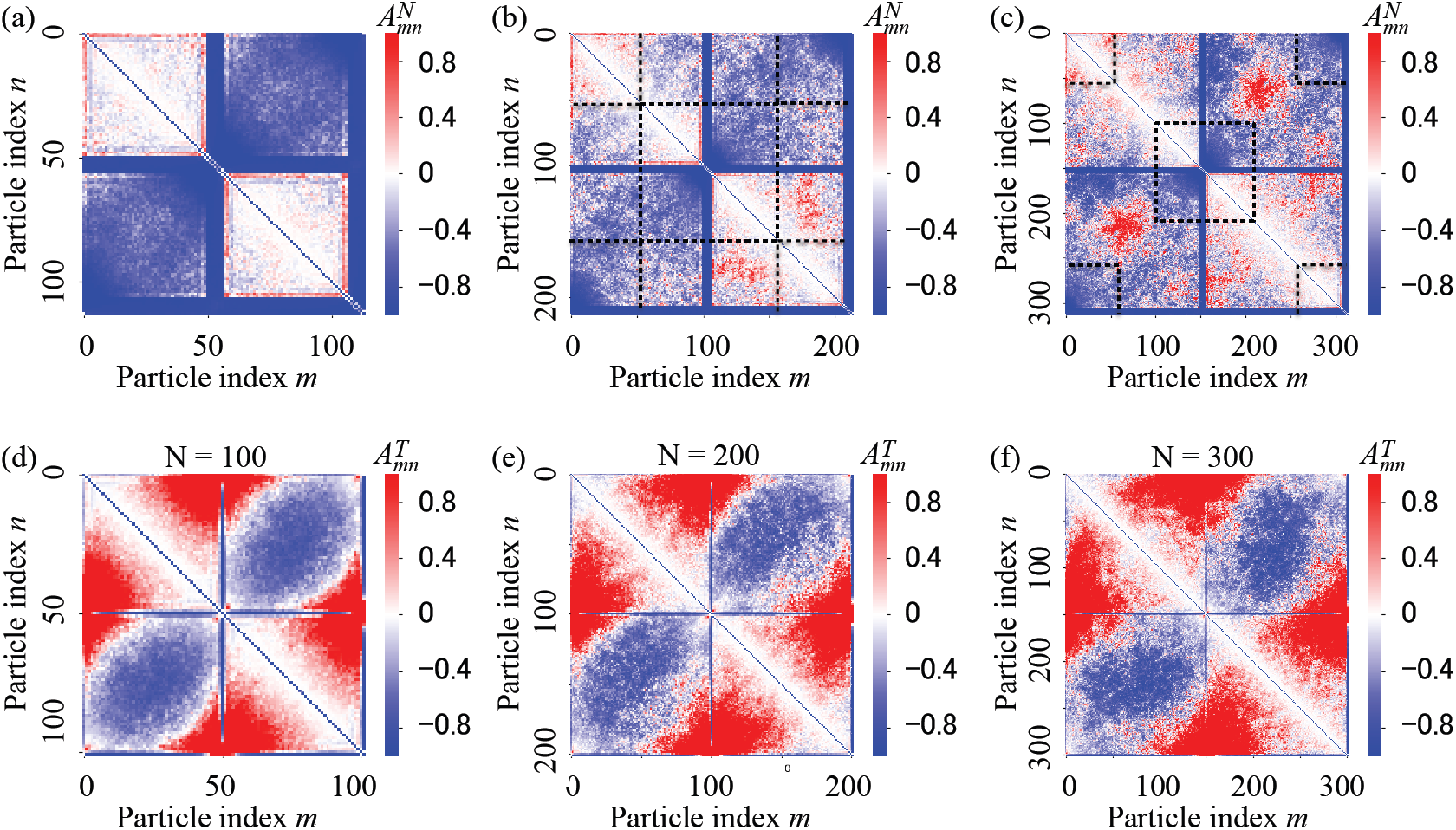
Relative contact probability matrices of NENLIS-ring and Two-coupled-ring models. 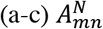 of NENLIS-ring for *N =* (a) 112 (approximately 18 kbp), (b) 212 (approximately 36 kbp), and (c) 312 (approximately 54 kbp), and 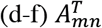 of Two-coupled-rings for *N =* (d) 100, (e) 200, and (f) 300 with *L* = 1.01*d* and *k*^*at*^*d*^2^/*D* = 200. NENLIS locates at particle indices (a) 50–55 and 106–111, (b) 100–105 and 206–211, and (c) 150–155 and 306–311. The distances of particles with indices *m* and *n* in dashed square areas in (b) and (c) from NENLIS along the particle chain were shorter than the length of 50 particles (approximately 9 kbp). Two particles with indices (d) 49 and 99, (e) 99 and 199, and (f) 149 and 299 were connected to permanently form two loops. Positive 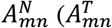) means that associations between the *m*-th and *n*-th particles occur more frequently in the NENLIS-loop (Two-coupled rings) than in the control ring, and vice versa.

We found that the majority of 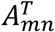 among the loci belonging to different loops were negative whereas those near boundary particles were positive independent of *N*, for *N* = 100, 200, and 300 (Figure 3d-f). Thus, the interactions among loci in different chromatin loops far from the chromatin domain boundary tended to be suppressed whereas those near the boundary tended to be enhanced. These results were consistent with recent studies of loop-forming insulators.^29,32^ Together, these findings indicated that loop-forming insulator sequences could function as a boundary between larger chromatin domains than those distinguished by NENLIS. However, for chromatin domains of around 10 kbp, which is longer than the frequently observed promoter-enhancer distances, NENLIS appeared to exhibit more accurate insulation than that provided by the loop-forming insulator sequences.

A recent study of Two-coupled-rings suggested that inter-nucleosome attractions were also important to realize the insulation between two chromatin domains ^42^. We also found that NENLIS did not strongly hinder the association between loci belonging to different nucleosome-rich regions in the NENLIS-ring in the case of small *k*^*at*^*d*^2^/*D* (Figure 4). This further indicated that inter-nucleosome attraction also played an important role in the formation of discrete chromatin domains.

**Figure 4.**
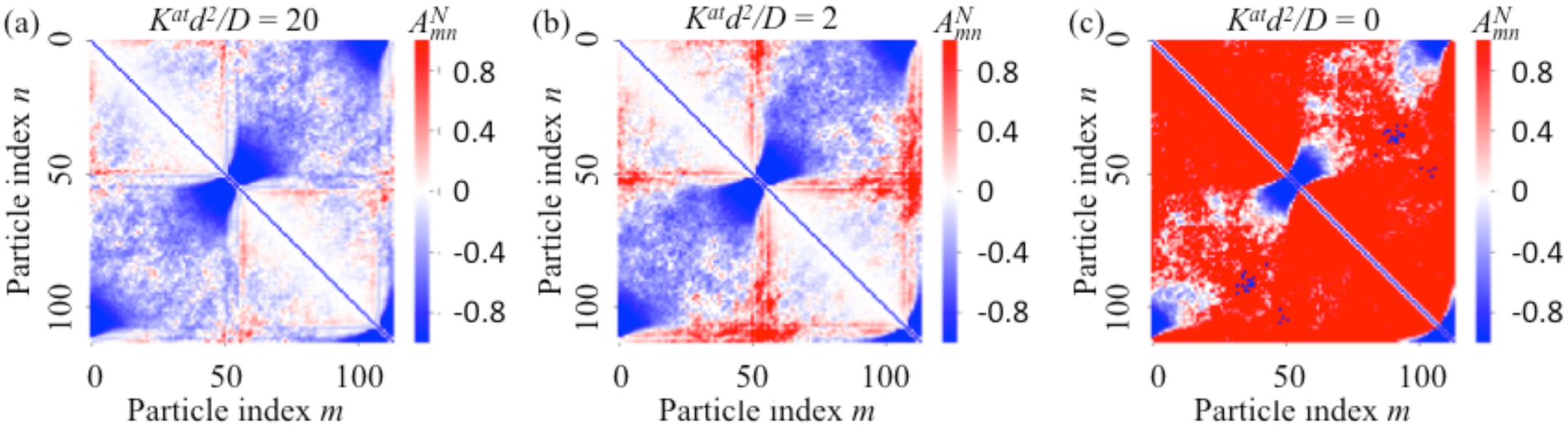
Relative contact probability matrices of the NENLIS ring with various inter-nucleosome interaction strengths. 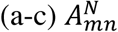 of the NENLIS-ring for *N =* 112 (approximately 18 kbp) with (a) *k*^*at*^*d*^2^/*D* = 20, (b) *k*^*at*^*d*^2^/*D* = 2, and (c) *k*^*at*^ = 0. NENLIS locates at particle indices 50–55 and 106–111.

### Expected Insulator Activities of NENLISc in Human GM12878 Cells

In human GM12878 cells, we identified 6812 sequences as NENLISc. We measured 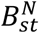 and 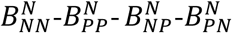 profiles for these (typical profiles are shown in Figure 5) and found that 5607 (around 82%) NENLISc could be considered as exhibiting insulator activities. These results suggested the possibility that the majority of NENLISc in the human genome have a similar role as NENLIS. We also found that the average associations among loci belonging to the same chromatin regions, 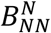, and 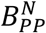, showed both positive and negative values (sometimes 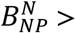 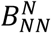 or 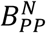 occured) depending on the NENLISc. This characteristic appeared to markedly differ from that of chromatin loop-forming insulator sequences, for which the loci belonging to the same chromatin regions were considered to associate with sufficient frequency to support the formation of topologically associated domains.

**Figure 5.**
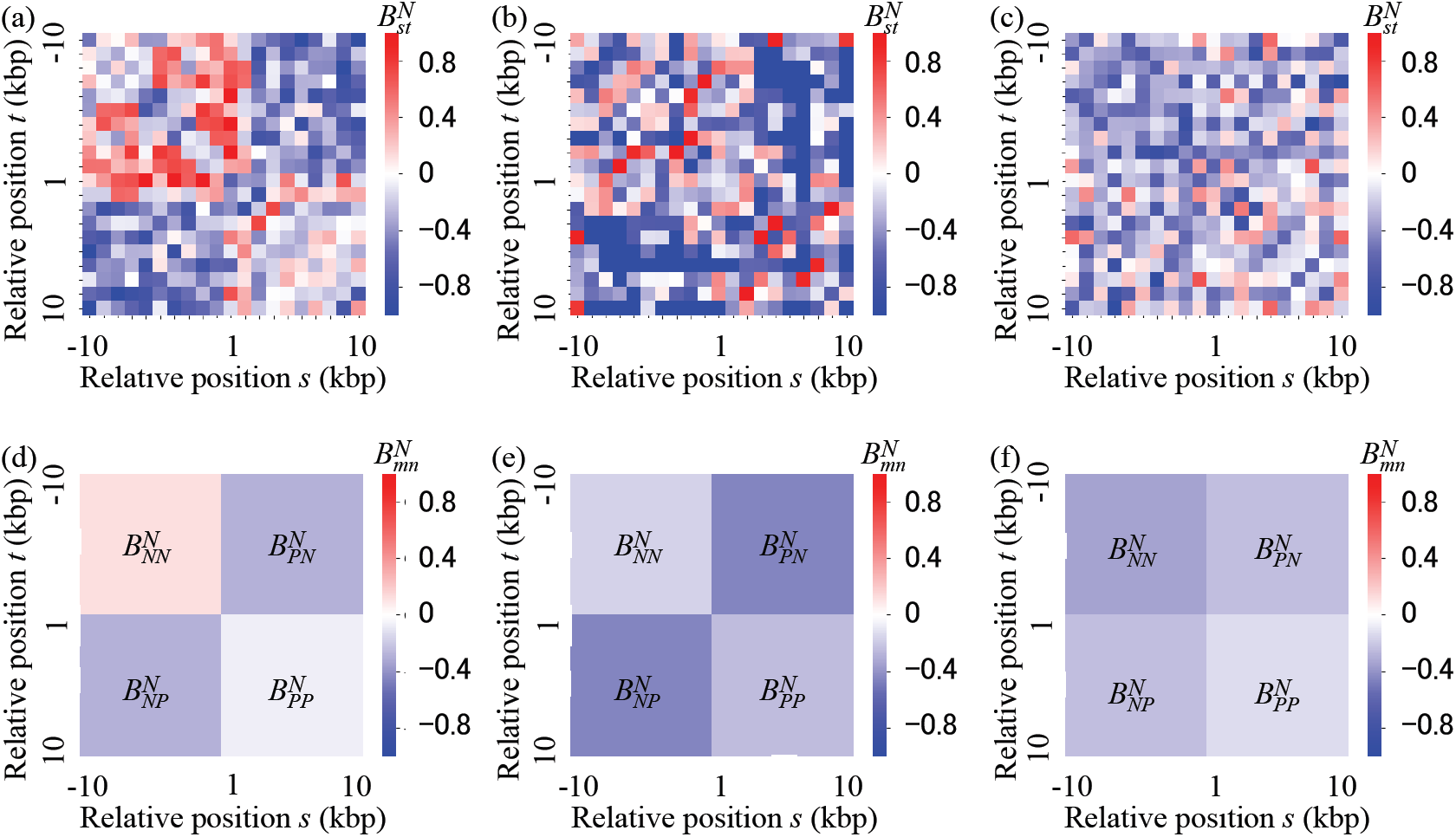
Relative contact probability matrices around NENLISc. (a-c) Typical 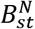 around NENLISc at loci (a) from the 33,365,515th to 33,365,667th bp in chr. 8, (b) 13,698,416th to 13,698,639th bp in chr. 7, and (c) 21,502,095th to 21,502,336th bp in chr. 13 in human GM12878 cells, and (d-f) 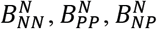 and 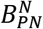 obtained from (a), (b), and (c), respectively. Relative position *s* and *t* indicates the relative genomic loci positions from NENLISc. 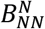 and 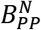 showed both positive and negative values as 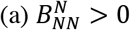 and 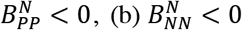 and 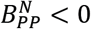, and 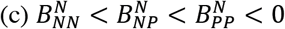 depending on the NENLISc, unlike those obtained by chromatin loop-forming insulator sequences.

Notably, various additional effects that were not considered in this argument should also be considered to more accurately assess the performance of NENLISc, such as histone modifications and protein binding. These represent important areas for future study.

## CONCLUSIONS

In this study, we investigated the insulator activities of NENLIS and NENLISc in the human genome using mathematical models and genome-wide analysis of Hi-C data of human GM12878 cells. First, we constructed a coarse-grained model of chromatin containing NENLIS wherein nucleosome-rich regions were expected to show relatively higher flexibility and could exhibit larger fluctuations upon long-term averaging than NENLIS owing to the activities of chromatin remodeling factors. Through simulations of this model we found that associations between two chromatin regions with approximately 10 kbp distance were sufficiently hindered by NENLIS. In addition, we also identified various NENLISc in GM12878 cells and showed that they could interfere with the associations between two flanking chromatin regions. These findings indicated that NENLIS could exhibit insulator activity only through nucleosome exclusion; moreover, such types of insulator sequences might exist widely in various organisms including human.

## AUTHOR INFORMATION

The authors declare no competing financial interest.

## ACKNOWLEDGMENT

This research was partly supported by the Platform Project for Support in the Japan Agency for Medical Research and Development (to A.A.); a Grant-in-Aid for Scientific Research on Innovative Areas “Initiative for High-Dimensional Data-Driven Science through Deepening of Sparse Modeling” from MEXT (no. 26120525 to A.A.); a MEXT KAKENHI grant (no. 17K05614 to A.A.); and a Grant-in-Aid for Scientific Research (C) (JSPS KAKENHI Grant Number JP17K07241 to N.S.). The authors thank H. Nishimori and T. Kameda for helpful discussions.

